# Much stronger coarse-to-fine visual processing in primate superior colliculus than primary visual cortex neurons

**DOI:** 10.64898/2026.06.23.734056

**Authors:** Yue Yu, Amarender R. Bogadhi, Matthias P. Baumann, Tatiana Malevich, Tong Zhang, Carlotta Trottenberg, Ziad M. Hafed

## Abstract

The visual system optimizes its signal processing properties to efficiently encode natural scenes. The superior colliculus (SC) and primary visual cortex (V1) both play important roles in visual-motor processing, and they both exhibit qualitatively similar visual responses. However, it is not clear whether the SC simply inherits V1’s efficient coding image processing optimizations or not, especially given that the SC receives a substantial amount of direct anatomical inputs from V1. Here, by performing matched experiments in the two brain areas, as well as with the same visual stimuli and in the same experimental animals, we show that the dynamics of coarse-to-fine visual image processing in the SC are much stronger than those in V1. In the SC, visual response latencies, being fastest for coarse patterns, are dictated by image spatial frequency, independently of the spatial frequency tuning curves of the neurons. On the other hand, V1 visual response latencies are largely dominated by visual response sensitivity, which is itself much more broadband than in the SC. These observations remain fundamentally unchanged in active vision gaze-shift scenarios eliciting visual reafferent responses in both brain areas. Our results suggest that coarse-to-fine visual image processing dynamics are most observable, and thus most relevant, in visually-responsive neurons driving foveating eye movements, like in the SC. Besides explaining behavioral evidence for saccadic facilitation by images that are consistent with the statistics of natural scenes, these results indicate that coarse-to-fine visual processing dynamics are much more of a collicular than a cortical visual phenomenon.

## Introduction

Active vision entails repetitive sampling of the sensory landscape via self-motion. To achieve such sampling, motor generation and control areas in the brain often also exhibit sensory sensitivity ^1–3^. However, the nature of such sensitivity has remained relatively unexplored for a long time. For example, even though it has been known for decades that the primate superior colliculus (SC) exhibits visual responses ^1,4^, which likely help in driving foveating eye movements ^5,6^, it was not always clear whether these visual responses are merely inherited from the primary visual cortex (V1), a primary source of collicular visual input ^7,8^, or whether they are functionally distinct. Indeed, SC visual responses are qualitatively similar to V1 visual responses, and both SC ^5,9,10^ and V1 ^11^ visual responses are correlated with saccade timing. Only recently was a direct comparison between the two brain areas made, in the same animals and with the same stimuli, to reveal a much larger than expected difference between the SC and V1 in terms of linking visual response properties to eye movement reaction times ^12,13^.

Among other visual parameters, an image processing domain that is particularly relevant for both scene analysis and scene exploration by gaze shifts is that of spatial frequency. Here again, functional specialization and efficient coding principles would anticipate that both V1 ^14^ and the SC ^15,16^ should be able to represent visual pattern information, which is indeed the case. However, the use of spatial frequency information for perception (a functional domain of V1) is not necessarily the same as its use for action (a functional domain of the SC). Consider, for example, the process of object recognition. Such recognition may be jumpstarted in peripheral vision by an initial detection process. Then, a foveating eye movement might ensue, to allow subsequent detailed visual inspection. In this case, peripheral object detection needs to be rapid, but it does not need to be exhaustive. Thus, low spatial frequency information may suffice for this critical first step, and this might explain why the oculomotor system, including the SC, exhibits express detection (but not necessarily recognition) capabilities for visual objects ^15,17–20^. On the other hand, if peripheral object recognition is needed without a foveating eye movement, then higher spatial frequency information becomes more relevant, which might require recruitment of V1 machinery. This example demonstrates that the temporal aspect of visual responses ^21^, whether in the SC or V1, is particularly relevant in pattern vision, explaining phenomena like coarse-to-fine image processing dynamics ^22–26^.

Here, we hypothesized that even though coarse-to-fine image processing dynamics exist in both V1 ^22–25^ and the SC ^15^, they would still exhibit large and fundamental differences across the two brain areas, exactly by virtue of the above naturalistic active vision scenario of early/coarse peripheral detection followed by more detailed analysis through foveation. Consistent with this, we found a particularly strong instantiation of coarse-to-fine dynamics, in the response latencies of visual responses, which was almost exclusively unique to the SC for the ranges of spatial frequencies that we focused on, and essentially absent in V1.

## Results

We explored the dynamics of coarse-to-fine visual image processing in the SC ^15,27^ and V1 ^22–25^ of the same animals, and with the same stimuli. Two rhesus macaque monkeys fixated a small spot, and we presented a static Gabor grating of a given spatial frequency within the receptive fields (RF’s) of SC and/or V1 neurons (Methods). We noted a dramatic difference in the temporal dynamics and response gain properties of coarse-to-fine image processing across the two brain areas. Most strikingly, there was a rank ordering of SC visual response latencies as a function of the presented spatial frequency, which was independent of neuron preference, and which was also virtually absent in V1.

### Much more delayed SC than V1 visual responses to high spatial frequencies

Figure 1A shows the visual responses of an example SC neuron to four different spatial frequencies. In terms of visual response gain, the neuron preferred (or was tuned to) 4 cycles/deg (cpd): it emitted the strongest visual response for this spatial frequency, and all other spatial frequencies elicited weaker responses. In terms of visual response latency, however, the neuron preferred 0.5 cpd, for which it responded the earliest. The top-right and bottom-right panels of Fig. 1A illustrate this dissociation between response gain and response timing in this neuron: the earliest visual response was not the strongest one, and this replicates our earlier observations obtained from the SC of two additional animals ^15^. A similar conclusion could be reached for the second example SC neuron of Fig. 1B. This time, the neuron preferred 1 cpd, with visual responses being weaker for both lower (0.5 cpd) and higher (2 cpd and 4 cpd) spatial frequencies (top-right panel). Nonetheless, the same rank ordering of response latencies with spatial frequency was observed as for the first example neuron of Fig. 1A (compare the bottom-right panels of both neurons).

**Figure 1.**
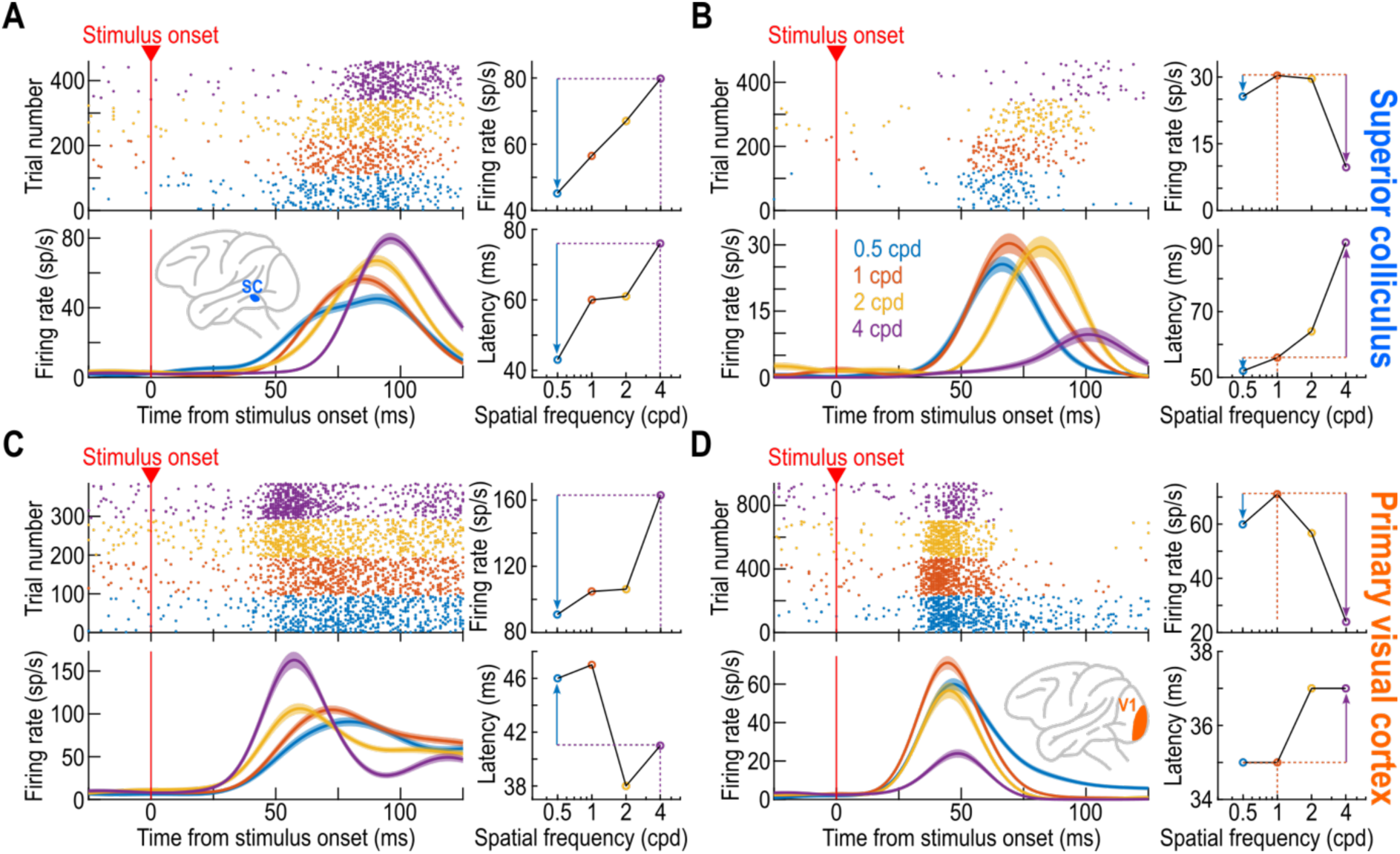
Delayed superior colliculus (SC), but not primary visual cortex (V1), visual responses for higher spatial frequencies, independent of visual response sensitivity. **(A)** Example SC neuron preferring 4 cpd in terms of visual response strength. The left two panels show the raw responses of the neuron (top: spike time rasters, sorted by spatial frequency; bottom: average firing rate curves across trials). The top right panel summarizes the visual sensitivity tuning curve of the neuron: peak visual response (Methods) was strongest for 4 cpd. Despite that, the bottom right panel shows that the visual response latency (Methods) was longest for the preferred spatial frequency, and it was shortest for the lowest spatial frequency ^15^. **(B)** Another example SC neuron, now preferring 1 cpd. The response latency for the lower spatial frequency (0.5 cpd) was still earlier than the response latency for the preferred spatial frequency (1 cpd), whereas the response latency for higher spatial frequencies (e.g. 4 cpd) was longer; thus, the neuron’s response latency favored low spatial frequencies irrespective of visual sensitivity ^15^; note how the response latency curve was qualitatively similar to that of A despite the difference in visual sensitivity curves. **(C)** An example V1 neuron preferring 4 cpd (like in A). Here, the response latency for 4 cpd was shorter than for 0.5 cpd, reflecting the neuron’s visual sensitivity. Thus, in V1, visual response latency was more dominated by the strength of the visual response ^21,30,31^. **(D)** An additional example V1 neuron, now preferring 1 cpd (like in **B**). Once again, the neuron’s visual response latency was governed more strongly by its visual response sensitivity than by the presented spatial frequency. Also see Fig. S1 for additional example neurons. Error bars denote SEM across trials, and trial numbers can be inferred from the spike raster panels.

The above observations were categorically different in V1. Consider, the example neuron of Fig. 1C. Like the SC neuron of Fig. 1A, this V1 neuron preferred 4 cpd. However, in this case, the response to 4 cpd was much earlier than the response to 0.5 cpd. In fact, the earliest visual response of this neuron was for a spatial frequency that was closest to its preferred one, consistent with the expectation that stronger visual responses normally also have shorter latencies ^5,12,21,28–31^. Similarly, the V1 neuron of Fig. 1D preferred 1 cpd (like the SC neuron of Fig. 1B). Once again, the earliest visual response was for the stimulus eliciting the strongest visual response, and not for 0.5 cpd as in the SC neurons. Four additional example SC and V1 neurons are shown in Fig. S1, and they all indicate that coarse-to-fine temporal dynamics were much stronger in the SC than in V1.

Across the population, we estimated the average visual response latency of each neuron for each spatial frequency (Methods). We then plotted cumulative distributions across all neurons. In the SC (Fig. 2A; top panel), we replicated our earlier findings ^15^, but with two additional monkeys that were different from those that were used in the previous study: there was a clear rank ordering of SC visual response latencies with increasing spatial frequency. In contrast, the V1 population of the same two animals as in the top panel of Fig. 2A had much weaker coarse-to-fine visual response temporal sequencing (Fig. 2A; bottom panel). In fact, the population visual response to 1 cpd in V1 was slightly earlier than that to 0.5 cpd (also see Fig. 2B). And, the response to 2 cpd was as early as that to 0.5 cpd; only for 4 cpd was there evidence for a delaying of V1 visual responses relative to the lower spatial frequencies. As shown in Fig. 2B, this delaying was only 11.2% relative to the visual response latency for 0.5 cpd, but it was more than twice as large (24.8%) in the SC (Fig. 2B). In units of absolute time, visual response latency for 4 cpd in the SC was ∼14 ms longer than for 0.5 cpd, but it was only ∼5 ms longer in V1. We also confirmed these observations statistically. A mixed-effects model applied to visual response latencies revealed main effects of brain area (F=69.74, p<0.0001, dF=1, n=363 neurons across areas) and spatial frequency (F=117.14, p<0.0001, dF=3). Moreover, there was an interaction between brain area and spatial frequency (F=19.507, p<0.0001, dF=3), consistent with the observation of a larger impact of the latter on visual response latencies in the SC than in V1. Thus, in terms of visual response latencies, the SC exhibited much stronger coarse-to-fine temporal dynamics than V1.

**Figure 2.**
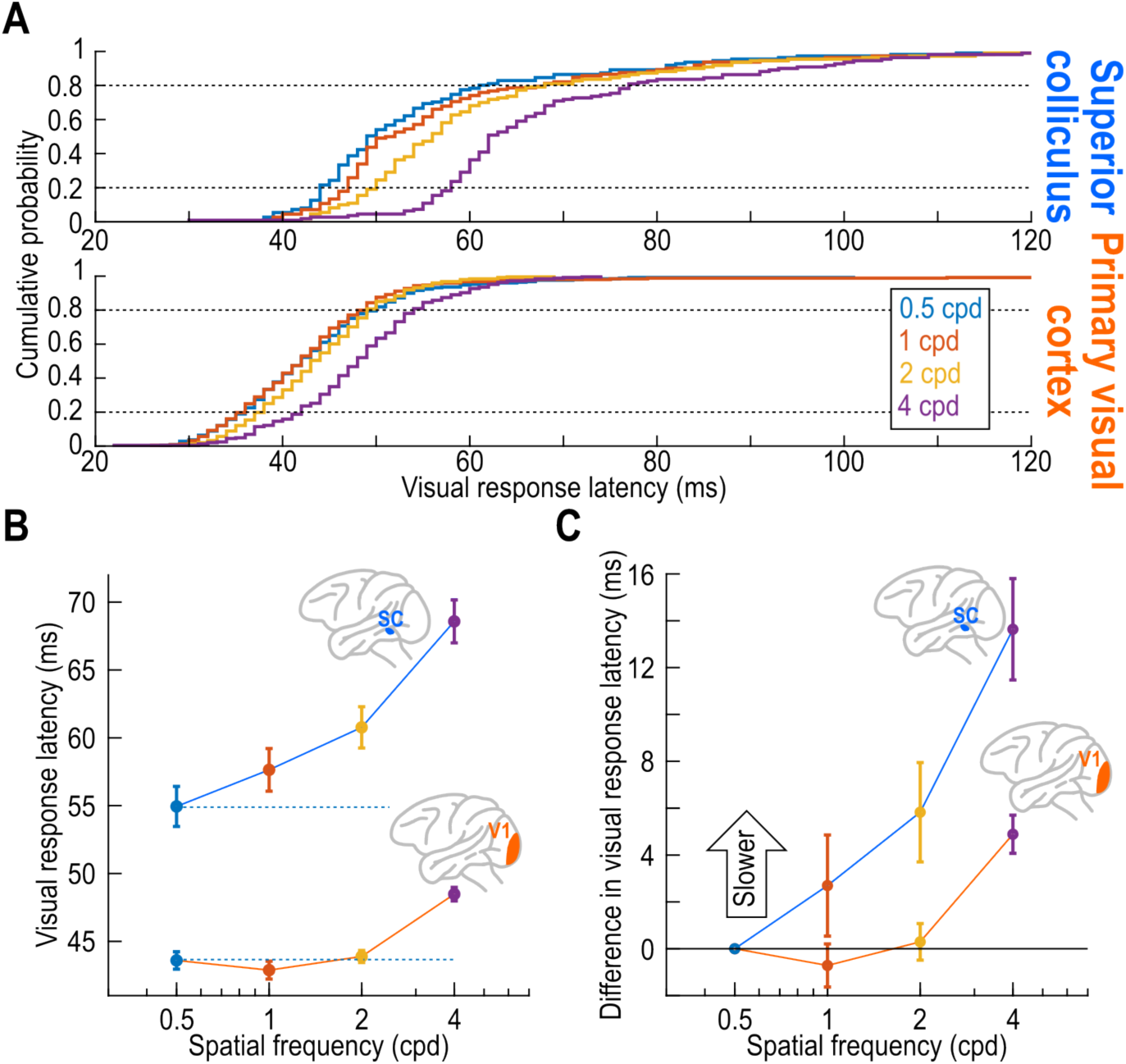
Much stronger coarse-to-fine visual processing in the primate SC than V1. **(A)** Each panel shows the cumulative histogram of visual response latencies of SC (top; n=112 neurons) and V1 (bottom; n=251 neurons) neurons in our database. Each curve shows the visual response latency distribution for a given spatial frequency. In the SC, there was a rank ordering of visual response latencies as a function of increasing spatial frequency ^15^. In V1, this effect was much weaker; in fact, the distribution for 1 cpd was slightly earlier than that for 0.5 cpd (also see **B**, **C**). **(B)** Summary of the results of **A**. Each symbol indicates the average visual response latency of the SC or V1 population for a given spatial frequency; error bars indicate SEM across neurons. In the SC, visual response latency increased with increasing spatial frequency (the dashed horizontal line indicates the response latency of the population for 0.5 cpd); for V1, this effect was much weaker. For example, the visual response latency at 4 cpd was 11.2% slower in V1 than the visual response latency at 0.5 cpd (48.48 ms versus 43.59 ms); in the SC, the visual response latency at 4 cpd was 24.8% slower than for 0.5 cpd (68.58 ms versus 54.95 ms). Note also that V1 visual response latencies were generally smaller than in the SC, consistent with our earlier direct latency comparisons within the same animals ^12^. **(C)** The data in **B** but now plotted as a difference from the average population visual response latency for 0.5 cpd spatial frequency. There was more systematic delaying of SC visual responses with increasing spatial frequency than in V1. Error bars denote SEM across neurons.

### *Dissociation between response sensitivity and response latency in the SC, but* not V1

Our previous studies revealed that dissociations between visual response sensitivity and visual response latency (Figs. 1A, B, S1A, B) could be routinely observed in the SC, not just in the context of spatial frequency ^15^, but also in other sensory dimensions, like luminance polarity ^32^. This suggests that such dissociations might be substantially stronger in the SC than in the cortex, but the results of Fig. 2 above were, so far, agnostic of response sensitivity. When we explored such sensitivity further, we confirmed that there was an almost-complete lack of dissociation between sensitivity and latency in V1, unlike in the SC.

To illustrate this, consider first the results of Fig. 3. Here, we picked all SC or V1 neurons that were tuned to one spatial frequency (1 cpd by way of example), and we compared visual response latency and visual response sensitivity for this spatial frequency to response latency and sensitivity for the other nonpreferred spatial frequencies. In both the SC (Fig. 3A, B) and V1 (Fig. 3E, F) response strength was higher for 1 cpd than for lower (0.5 cpd) and higher (2cpd, Fig. 3A, E; 4 cpd, Fig. 3B, F) spatial frequencies; this is expected from our a priori selection of only the neurons preferring 1 cpd. However, when analyzing response latencies for the same neurons, the SC and V1 populations were again categorically different from each other. In the SC, the population response latency for 0.5 cpd was consistently earlier than the response latency for 1 cpd even though the neurons preferred 1 cpd (blue population of neurons in Fig. 3C, D). On the other hand, the population response latency for 2 cpd (Fig. 3C) and 4 cpd (Fig. 3D) was later than the response latency for 1 cpd: in other words, note how the neurons were systematically on opposite sides of the unity-slope diagonal line for the different spatial frequencies in Fig. 3C, D, even though they were on the same side of it in terms of response strength (Fig. 3A, B). Thus, the response latencies of the SC neurons were dictated by the presented spatial frequency and not by the neurons’ visual sensitivity. In stark contrast, the V1 neuron response latencies were not faster for the nonpreferred spatial frequencies than they were for the preferred (1 cpd) spatial frequency (Fig. 3G, H; all statistical tests are included in Fig. 3). Therefore, in V1, visual response latencies were primarily dictated by visual response sensitivity; there was no strong V1 dissociation between visual response latency and visual response sensitivity like there was in the SC.

**Figure 3.**
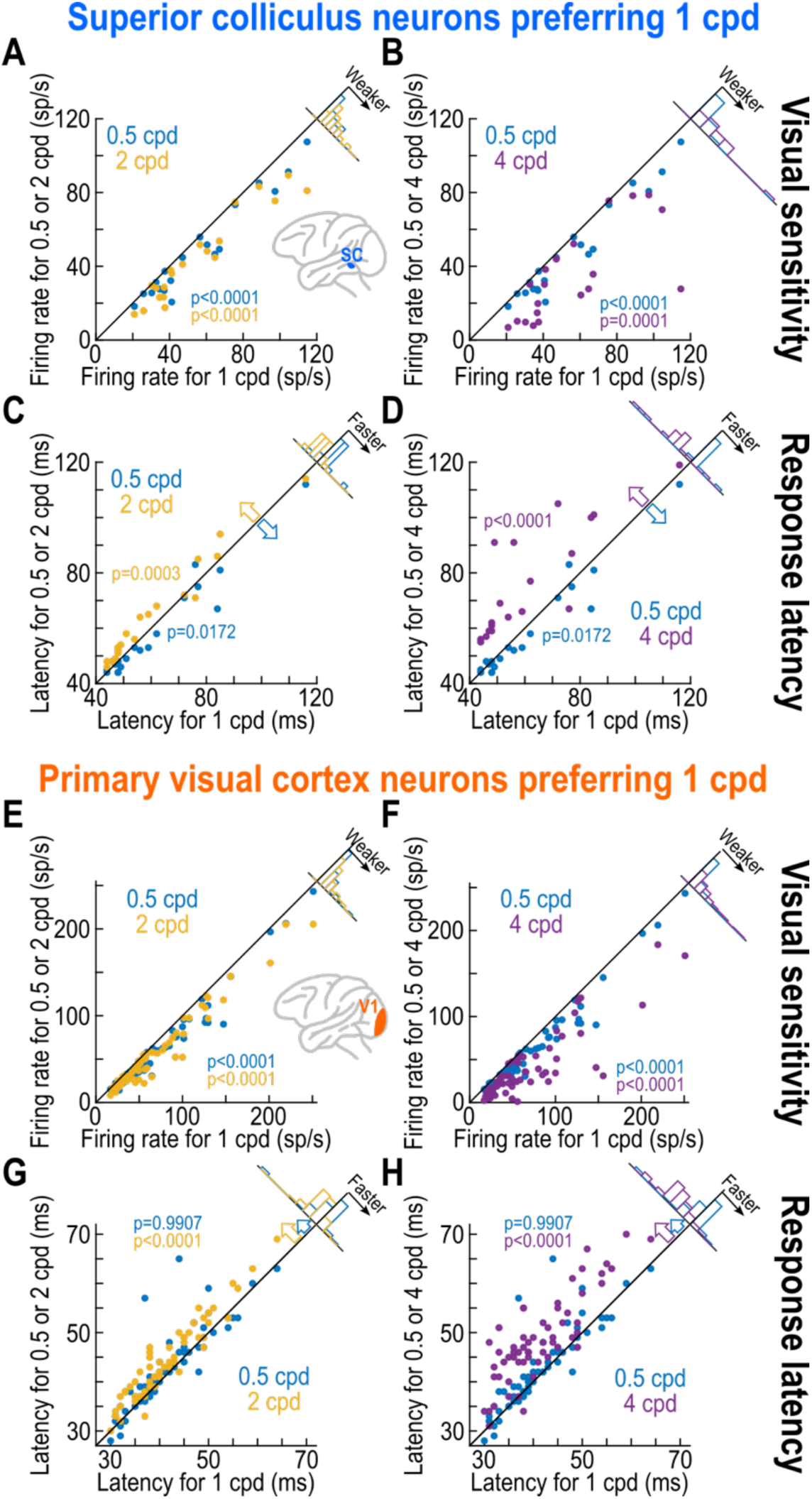
Dissociation between visual response strength and visual response latency in the SC, but not V1. **(A)** For all SC neurons preferring 1 cpd, we plotted the visual response strength of the neurons for either 0.5 cpd (lower than 1 cpd) or 2 cpd (higher than 1 cpd) against the visual response strength for 1 cpd. As expected from the selection of neurons preferring 1 cpd, visual response strength was weaker for both 0.5 cpd (blue) and 2 cpd (yellow) than for 1 cpd. P-values indicate the results of pairwise t-tests (corrected for multiple comparisons; Methods) for each of the shown analyses. For the marginal distributions, there was also no significant difference (t-test, p=0.2187, n=19), suggesting similar decreases in response strength for the two nonpreferred spatial frequencies. **(B)** Similar observations when the higher spatial frequency was 4 cpd (purple) instead of 2 cpd. For the marginal distributions, there was a significant difference (t-test, p=0.0110, n=19), indicating a bigger difference between 4 cpd and 1 cpd than between 0.5 cpd and 1 cpd. **(C)** For the same comparison as in **A**, we now plotted visual response latency (Methods). Even though the neurons preferred 1 cpd, their visual responses for the lower spatial frequency (0.5 cpd; blue) came earlier than they did for the preferred spatial frequency. For the higher spatial frequency (2 cpd; yellow), the visual responses arrived later. As a result, the marginal distributions were also statistically significantly different from each other (t-test, p=0.0001, n=19). Thus, in the SC, there were earlier visual responses for low spatial frequencies even when they were not preferred ^15^. **(D)** Similar observations when including an even higher spatial frequency relative to the preferred one. The marginal distributions were again different from each other (t-test, p<0.0001, n=19). **(E-H)** Same as **A**-**D** but for V1. In this case, the visual response latency for 0.5 cpd (**G**, **H**; blue) was not faster than the visual response latency of the preferred 1 cpd. For the higher nonpreferred spatial frequencies (2 cpd in **G** and 4 cpd in **H**), the visual response latencies were longer, consistent with the weaker responses. Thus, in V1, visual response latencies were more governed by visual response strength. The statistics comparing marginal histograms in each panel were as follows: p=0.2988, n=61 for **E**; p<0.0001, n=61 for **F**; p=0.0289, n=61 for **G**; p<0.0001, n=61 for **H**.

We also confirmed these observations for the populations of SC and V1 neurons preferring other spatial frequencies. First, we plotted normalized average population firing rates for each spatial frequency, but after separating the neurons according to their spatial frequency preference ^15^ (Methods). In the SC (Fig. 4A), the rank ordering of population firing rate curves with increasing spatial frequency was maintained for each subpopulation of SC neurons, just like in our earlier report ^15^. Thus, whether a neuronal population preferred 0.5 cpd (leftmost panel in Fig. 4A) or 4 cpd (rightmost panel of Fig. 4A), the visual responses arrived earlier for 0.5 cpd than for 4 cpd (colored oblique arrows in all panels of Fig. 4A) ^15^. Once again, this observation was qualitatively, not merely quantitatively, altered in V1 (Fig. 4B); in this case, the strongest visual response (for the preferred spatial frequency) was also generally the earliest visual response (colored oblique arrows in Fig. 4B).

**Figure 4.**
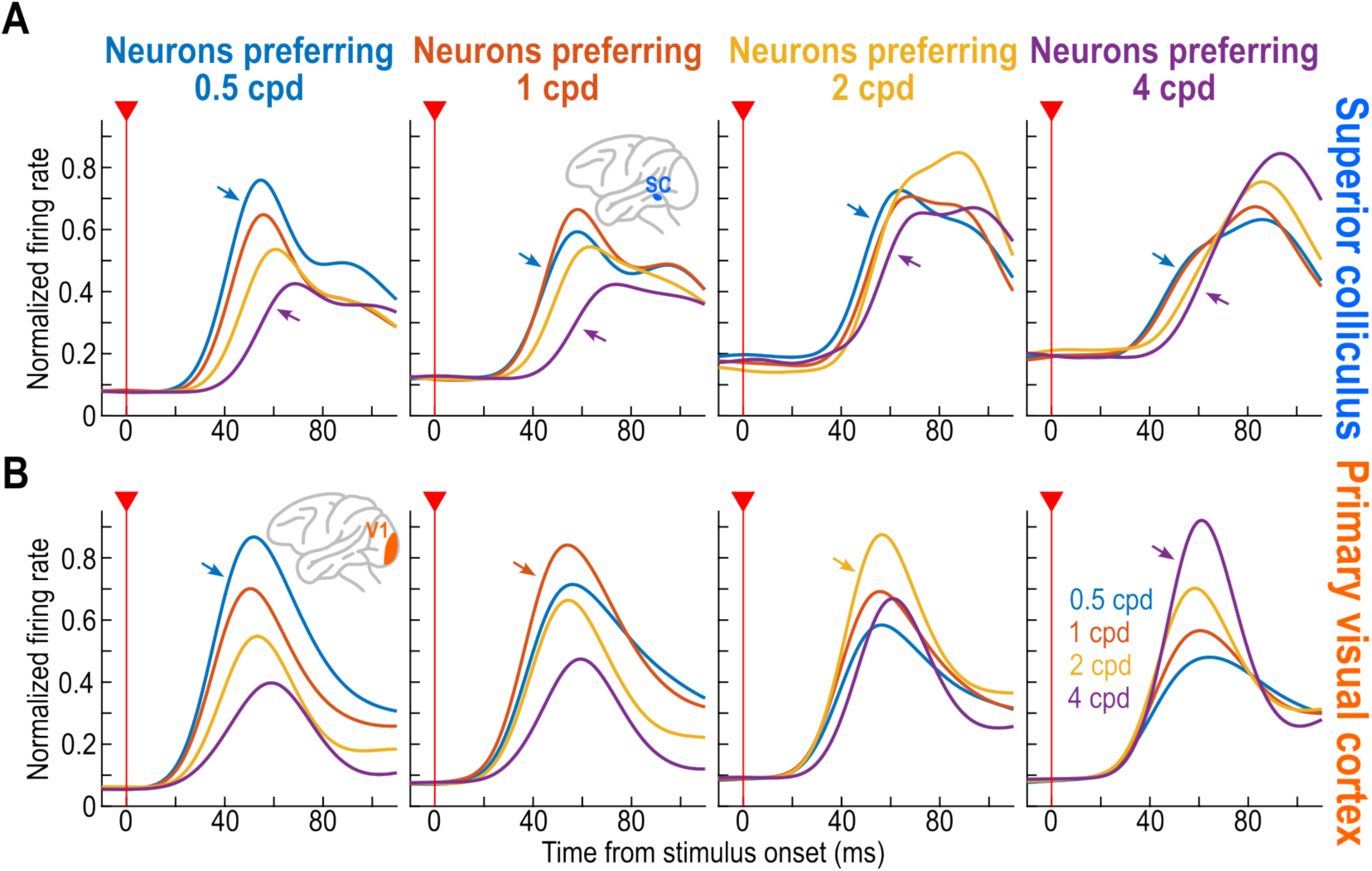
Rank ordering of SC, but not V1, visual response latencies with spatial frequency, independent of response strength. **(A)** For neurons preferring each of the presented spatial frequencies (each column), we plotted the average normalized firing rate of the SC population for each spatial frequency (Methods). Neurons preferring 0.5 cpd had the highest visual response for 0.5 cpd (leftmost column), whereas neurons preferring 4 cpd had the highest visual response for 4 cpd (rightmost column). For each subpopulation of SC neurons, the visual response for the lowest spatial frequencies (e.g. 0.5 cpd, small blue oblique arrows) appeared earlier than for the higher spatial frequencies (e.g. 4 cpd, small purple oblique arrows). Thus, the dissociation seen in Fig. 3 persisted for neurons preferring spatial frequencies other than 1 cpd (also see Fig. 5). **(B)** In the case of V1, the visual response latency for the preferred spatial frequency of each V1 subpopulation tended to be the lowest (small colored oblique arrows). Thus, in V1, visual response latency was more governed by visual response strength than by the presented spatial frequency.

We also repeated the analyses of Fig. 2, but for each subpopulation of neurons according to their preferred spatial frequency. In Fig. 5A, each curve shows the relationship between SC visual response latency and the presented spatial frequency for one subpopulation of neurons preferring one spatial frequency. Thus, the blue curve shows the visual response latencies of all SC neurons preferring 0.5 cpd, whereas the purple curve shows the visual response latencies of all SC neurons preferring 4 cpd (and similarly for the two other spatial frequencies). In each subpopulation of neurons, the image eliciting the fastest visual response was always the lowest spatial frequency (colored oblique arrows in Fig. 5A), even though this image was not preferred in all but the blue curve in the figure. The net result is that the relative delaying of SC visual responses with higher spatial frequencies (relative to 0.5 cpd) was maintained regardless of neuronal preference (Fig. 5B). On the other hand, in V1, the earliest visual response shifted towards higher spatial frequencies when the neurons preferred such higher spatial frequencies (colored oblique arrows in Fig. 5C). The net result was that when a neuron preferred a spatial frequency higher than 0.5 cpd, its visual response latencies for such spatial frequencies could be shorter than its visual response latencies for 0.5 cpd (Fig. 5D). Only for 4 cpd stimuli was there evidence of further delaying of V1 visual responses that was not explained by neuronal preferences (for example, there was no further decrease in response latency for the purple curve beyond 2 cpd, as might be expected if the preference for 4 cpd was the sole determinant of the response latencies at 4 cpd). Statistically, spatial frequency preference had no significant effect on SC response latencies (mixed-effects model, F=1.8213, p=0.1426, dF=3, n=112 neurons), but it did have a significant effect on V1 response latencies (mixed-effects model, F=18.35, p<0.0001, dF=3, n=251 neurons). Moreover, there was an interaction between spatial frequency and neuronal preference in V1 (mixed-effects model, F=9.3093, p<0.0001, dF=9, n=251).

**Figure 5.**
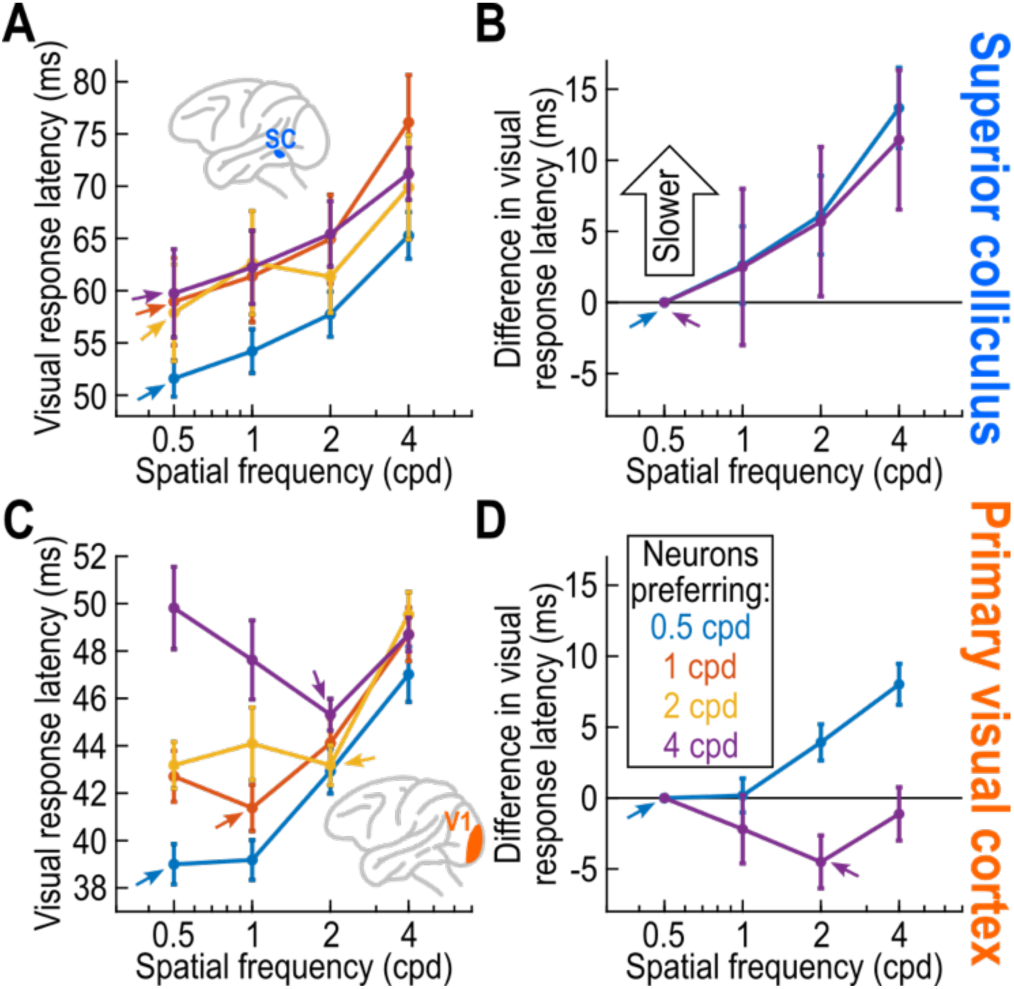
Dominance of coarse-to-fine visual processing in the SC. **(A)** Similar to Fig. 2, except that we first divided the SC population according to which spatial frequency was preferred. For the subpopulation preferring 0.5 cpd (blue curve), the shortest visual response latency was for 0.5 cpd spatial frequencies. This effect was not due to the strength of the visual response because 0.5 cpd was also associated with the shortest visual response latencies for each of the other three subpopulations of SC neurons preferring higher spatial frequencies (colored oblique arrows indicate the condition with the shortest visual response latency for each SC subpopulation). **(B)** Similar to Fig. 2C for the SC subpopulations preferring 0.5 cpd (blue) and 4 cpd (purple). In both cases, the relative delaying of visual response latencies for higher spatial frequencies was similar, independent of neuron preference. Thus, coarse-to-fine processing in the SC dominated this structure’s visual response latency relationships. **(C)** For V1, the stimulus eliciting the shortest visual response latency was the stimulus closest to the preferred stimulus. For example, for neurons preferring 1 cpd (reddish curve), the visual response latencies of these neurons were shortest when the presented stimulus had 1 cpd spatial frequency (reddish oblique arrow). Similarly, for neurons preferring 0.5 cpd, the shortest visual latency was for 0.5 cpd (blue curve and oblique arrow). **(D)** Same as **B** but for the V1 subpopulations preferring 0.5 cpd and 4 cpd. For the former, the earliest responses were for 0.5 cpd; for the latter, visual response latencies were slowest for 0.5 cpd, unlike in the SC. Note how the curve for 4 cpd indicates that some aspect of coarse-to-fine processing did remain in V1 because the fastest visual response latency was for 2 cpd instead of the preferred 4 cpd. Error bars denote SEM across neurons.

Therefore, the results of Figs. 3-5, combined, suggest that visual response latencies in V1 were largely determined by visual response sensitivity, whereas visual response latencies in the SC were largely determined by the presented spatial frequency. There was a much stronger coarse-to-fine dynamic in SC than V1 visual responses.

### Emphasis of SC coarse image processing in terms of visual sensitivity

Besides response timing, SC spatial frequency tuning curves also favor low spatial frequencies in terms of response sensitivity ^15^. Here, we had the opportunity not only to confirm this observation, but to also directly compare SC and V1 spatial frequency tuning curves at matched visual eccentricities in the same animals (which was never done before in the literature). Across the population, SC neurons were much more low-pass than V1 neurons. For example, Fig. 6A shows the distribution of neuronal preferences for our four presented spatial frequencies. In the SC, at the eccentricities that we tested (Figs. S2A, S3), 54.46% of the neurons preferred the 0.5 cpd spatial frequency (Fig. 6A). In contrast, the V1 neuronal preferences were essentially uniformly distributed across all four spatial frequencies (Fig. 6B). Thus, the notion of much stronger coarse-to-fine visual image processing in the SC than in V1 extends to both response sensitivity and response latency.

**Figure 6.**
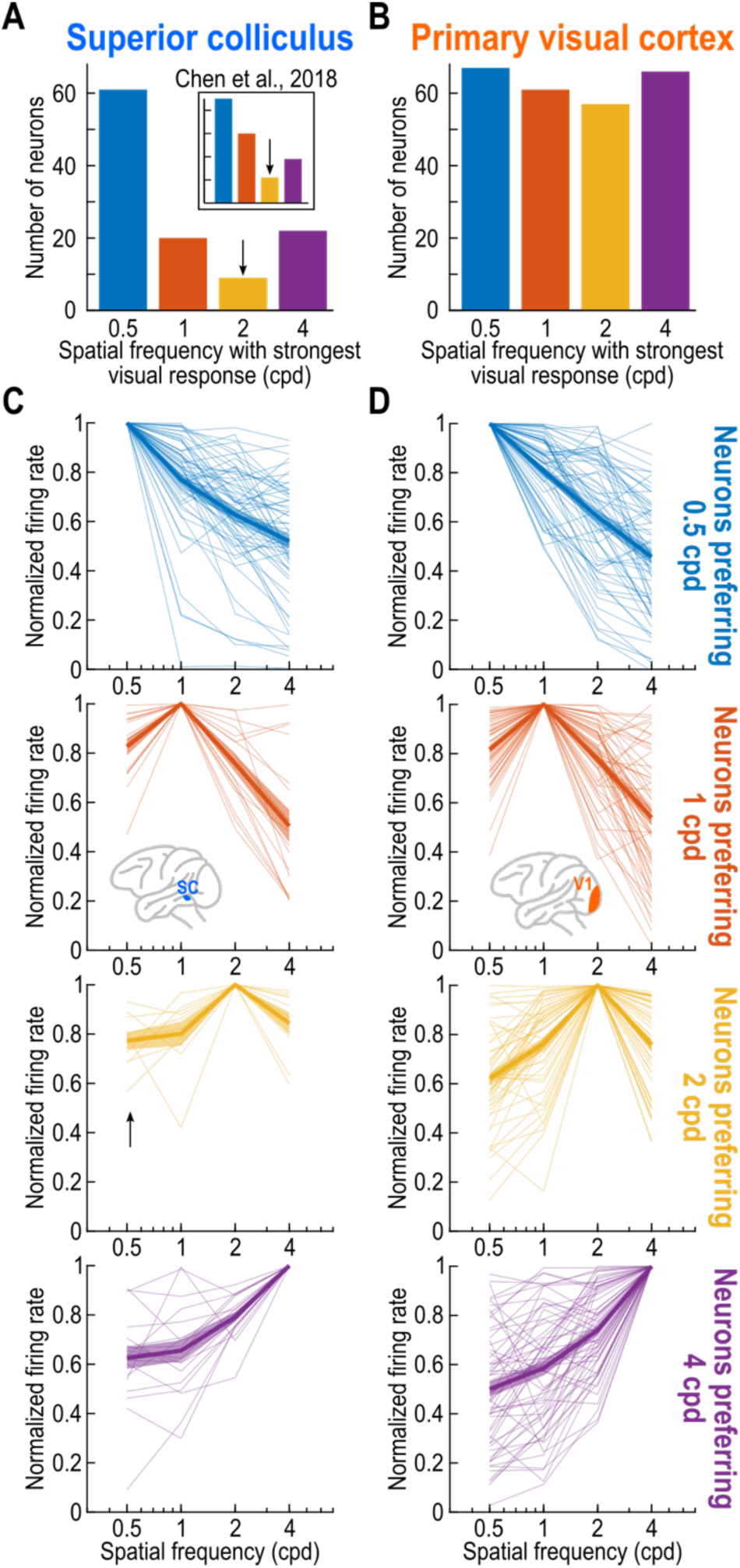
Emphasis of coarse image processing in the SC but not V1. **(A)** Distribution of preferred spatial frequencies in our SC population. Most neurons preferred 0.5 cpd. Note that there was a particular scarcity of SC neurons preferring 2 cpd (downward black arrow). This is likely a genuine property of the SC because inspection of the earlier report ^15^ with two additional animals revealed the same effect (the inset replicates the data from the earlier report for earlier comparison; note that the exact numerical values of the spatial frequencies were slightly different in the two studies, but they were similar in terms of range: 2.22 cpd as the highlighted spatial frequency in the inset versus 2 cpd in the current study). **(B)** V1 neuron preferences were uniformly distributed across all shown spatial frequencies. Also see Fig. S2 for a representation of the eccentricities that we tested, and Fig. S3 for how the spatial frequencies preferences appeared as a function of retinotopic eccentricity in our tested ranges. **(C)** Each row shows the population SC tuning curves for neurons preferring a given spatial frequency. For neurons preferring 2 cpd, the tuning curves were particularly shallow (black upward arrow), again suggesting a scarcity of preference for 2 cpd in the SC. **(D)** Same as **C** but for the V1 population. The tuning curve widths in the two areas were generally similar to each other, except for neurons preferring 2 cpd. Also see Fig. S4 for a more detailed comparison of tuning curve widths in the two areas. Error bars in **C**, **D** denote SEM across neurons. The inset of **A** was reproduced from ^15^.

Interestingly, in the SC, there was a particular scarcity of neurons preferring 2 cpd, relative to the other tested spatial frequencies (downward arrow in Fig. 6A). This scarcity was not a result of inevitable experimental variability in sampling neurons from the recorded brain structure, because it was remarkably also present in our earlier study (the data from that study are replicated in the inset of Fig. 6A for easier comparison) ^15^. Thus, an added advantage of replicating the SC experiments for our present purposes was that it allowed us to observe (and conclude), with an independent and larger neuronal database, that there exists a real and fundamental property of SC spatial frequency tuning curves: intermediate spatial frequencies (around 2 cpd) are particularly scarcely represented, unlike in V1.

We also had the opportunity to document spatial frequency tuning widths in the two brain areas. Figure 6C, D shows the population spatial frequency tuning curves of the two brain areas. In each row of these panels, we plotted the spatial frequency tuning curves of SC (Fig. 6C) or V1 (Fig. 6D) neurons preferring a given spatial frequency. Across brain areas, the tuning curve widths were generally similar to each other (also see Fig. S4), but neurons preferring 2 cpd were a clear exception: the SC tuning curves for this spatial frequency were particularly shallow, confirming the relative inability of the SC to represent 2 cpd spatial frequencies (alluded to in Fig. 6A). The net result is that across the entire population, SC tuning curves were rendered shallower than V1 tuning curves in the low frequency component of these tuning curves (Fig. S4C). This is unlikely due to lack of coverage by RF’s, especially because the population RF coverage relative to our stimulus sizes was more optimized for the SC neurons than for the V1 neurons (Fig. S2B, C; Methods).

Therefore, our results so far show a much stronger coarse-to-fine visual image processing machinery in the SC than in V1, whether in terms of visual response latency (Figs. 1-5), visual response sensitivity (Fig. 6A, B), or (at least partially) tuning curve widths (Fig. 6C, D).

### Similarity of results with visual reafferent responses after microsaccades

Finally, it was recently suggested that in mouse V1, there is a coarse-to-fine image processing dynamic, especially in active vision scenarios with saccadic gaze shifts ^33^. Because mouse V1 can be fundamentally different from primate V1, in both structure and function ^34–36^, the authors of that study also tested marmoset monkey V1. While they did confirm the mouse observations in the marmosets, these marmoset results were again obtained under active vision scenarios. Moreover, the experiments did not test the marmoset SC. Therefore, to ask whether our findings could potentially align with these earlier V1 observations from other species, we next considered SC and V1 visual reafferent responses after saccades.

In our experiments, the visual stimulus remained in the RF after its onset for a few hundred milliseconds (Methods). This allowed us to investigate visual reafferent responses of SC and V1 neurons after the occurrence of fixational microsaccades, like we previously did in the SC ^27^. We specifically investigated whether V1 neurons could exhibit stronger coarse-to-fine dynamics in these microsaccade-evoked visual reafferent responses than in the stimulus-evoked ones that we documented above.

Figure 7A, B shows the responses of two example V1 neurons when the stimuli were jittered in the neurons’ RF’s by microsaccades. In both cases, there were clear visual reafferent responses after the rapid eye movements. The neuron in Fig. 7A preferred 0.5 cpd in its reafferent response, and this condition also gave rise to the earliest reafferent visual burst timing (evident from the shown spike rasters). For the neuron in Fig. 7B, the preferred stimulus in terms of visual reafferent response strength was 2 cpd (when considering the peak of the firing rate curve), but the reafferent response for that spatial frequency was slightly delayed relative to the response for 0.5 cpd and 1 cpd spatial frequencies (again, evident from the shown spike rasters). Thus, this neuron could be considered as possessing a slightly clearer coarse-to-fine dynamic in its visual reafferent responses, particularly in the sense of a dissociation between sensitivity and latency like seen in the SC with our earlier stimulus-evoked analyses (Figs. 1, S1, 3-5).

**Figure 7.**
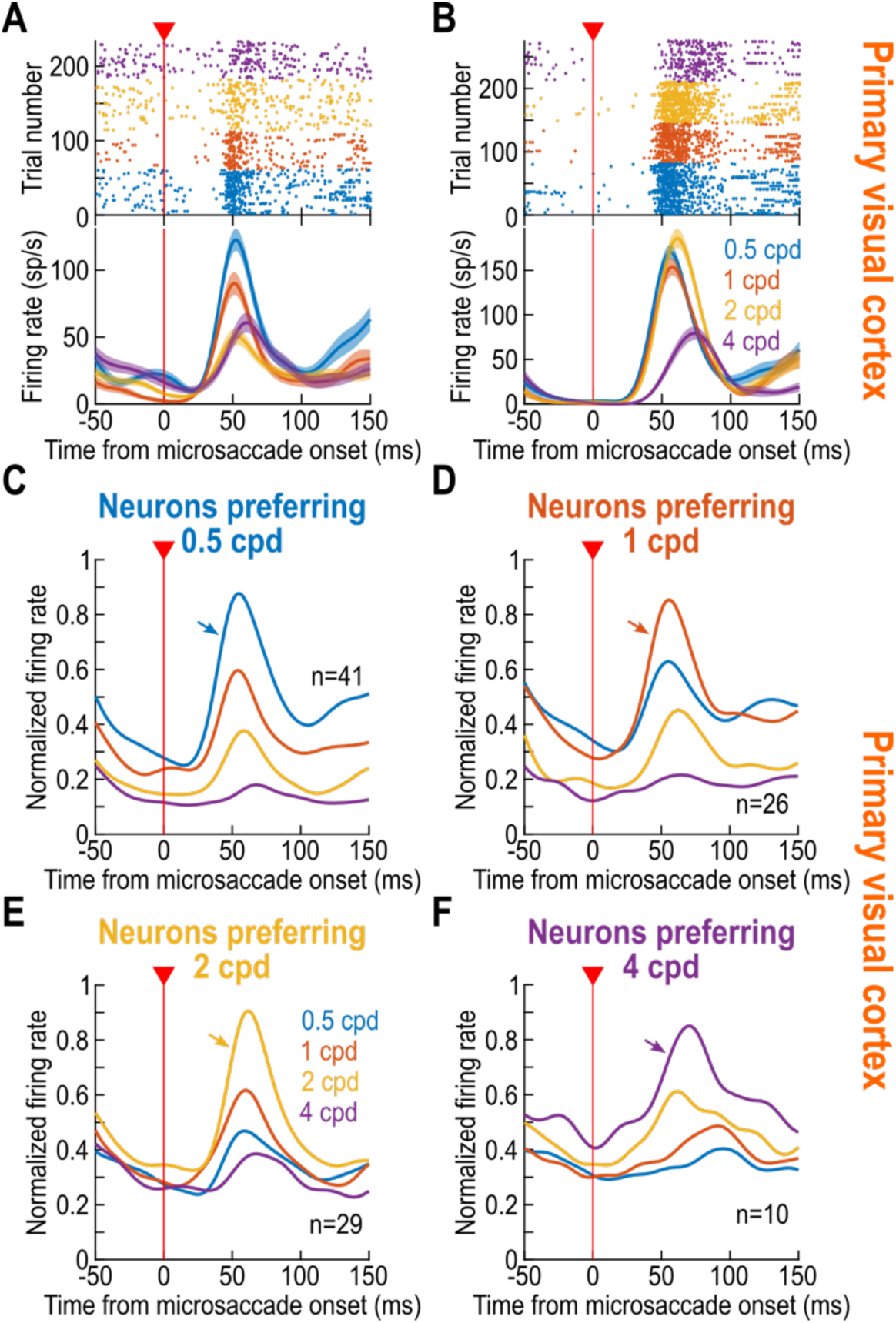
Similarity of V1 results with saccade-induced visual reafferent responses. **(A)** Responses of an example V1 neuron after a microsaccade has jittered the image of the presented visual stimulus over the neuron’s RF. The neuron had the strongest and earliest visual response for the lowest spatial frequencies. Error bars denote SEM across trials, and the number of trials can be inferred from the spike rasters. **(B)** Same as **A**, but for a second example V1 neuron. Now, the peak visual response was for 2 cpd. The latency of the visual reafferent response marginally slower for 2 cpd than the smaller spatial frequencies. **(C-F)** We grouped the V1 population as a function of which stimulus elicited the strongest visual reafferent response after microsaccades (Methods). In all cases, the average response latency was largely dominated by the response strength ^21,30,31^. Also see Figs. S5, S6.

Across the population of V1 neurons, however, V1 visual reafferent responses exhibited similar timing dependencies to those after stimulus onset. For example, Fig. 7C-F shows analyses similar to those shown in Fig. 4B. Specifically, we divided the V1 neurons according to which spatial frequency they preferred in their visual reafferent responses, and we then plotted the average normalized population firing rates for each group. Like in Fig. 4B, the preferred stimulus elicited the strongest and earliest reafferent responses. Indeed, when 0.5 cpd was preferred (Fig. 7C), its population reafferent response was earlier than for 4 cpd, but when 4 cpd was preferred (Fig. 7F), its population reafferent response was now much earlier than the response for 0.5 cpd. Therefore, once again, the timing of V1 visual reafferent responses was largely determined by visual reafferent response strength, and not by the presented spatial frequency.

Having said that, our V1 neurons did become slightly more biased to prefer lower spatial frequencies in their visual reafferent responses after microsaccades than with stimulus-evoked responses. This can be seen in Fig. S5. In this figure, the global population average firing rate after microsaccades was strongest for low spatial frequencies (Fig. S5A), and this was because most neurons now preferred the lowest spatial frequency in their reafferent responses (Fig. S5B). Thus, in this active vision context, our results in V1 could better align with the observations in other species ^33^, especially because in V1, stronger responses would also arrive earlier. Interestingly, the results of Fig. S5A, B suggest that the preferred spatial frequency of V1 neurons could be altered in the visual reafferent response epoch. This was indeed the case (Fig. S5C), and this could be due to interactions between the eye movement vector, its amplitude, and the orientation and spatial frequency of the stimulus, as we previously demonstrated in the case of the SC ^27^.

For completeness, we also analyzed the SC visual reafferent responses as well: we replicated our earlier findings ^27^, demonstrating once again an emphasis on coarse image processing, in both reafferent response timing and reafferent response sensitivity (Fig. S6).

## Discussion

We found a large difference between the primate SC and V1 in terms of visual pattern analysis capabilities. Besides being more sensitive to low spatial frequencies, the SC exhibits a dissociation between visual response latency and visual response sensitivity, which is much less obvious in V1 (at least in the ranges of spatial frequencies that we focused on). These observations could not be easily inferred from the prior literature, since there has been no direct comparison of the two brain areas in the same animals and with the same stimuli. Thus, it was difficult to assess whether prior evidence of V1 coarse-to-fine processing ^22–25^

was similar to, stronger than, or weaker than similar evidence in the SC ^15,27^. With our current comparison, we could now add to growing evidence that SC visual responses are not merely inherited from V1 ^1,12,13,15,28,32,37,38^. Rather, they are reformatted to reflect the functional contributions of the SC to active vision. In fact, visual signals in the SC are even still relevant at the time of saccade-related motor bursts, and in motor-dominated SC neurons ^38–45^. Such an integration of visual sensitivity into the SC matters a great deal for orienting behaviors. For example, much stronger SC coarse-to-fine image processing is a perfect substrate for saccadic facilitation in low spatial frequency scene contexts ^15,17,18^. Coupled with other visual asymmetries/dependencies in the SC, along with their established links to saccadic reaction times ^5,9,10,12,13,32,37,46^, these observations demonstrate the importance of visual responses in a classically-considered oculomotor control structure like the SC ^6,47,48^.

Besides foveating saccades, other oculomotor phenomena that are driven by sensory stimuli also exhibit clear coarse-to-fine dependencies in their strength and timing. For example, reflexive saccadic inhibition ^2,49–51^ is earlier and stronger for low spatial frequencies ^52^. Similarly, a related stimulus-driven ocular position drift phenomenon ^53,54^ is also stronger and earlier for low spatial frequencies ^53^. Interestingly, at least some of these oculomotor phenomena, such as saccadic inhibition, are critically dependent on V1 activity ^55^; however, despite this dependence, their feature tuning properties still reflect the feature tuning properties of visual signals in oculomotor circuits (such as the SC). This observation suggests that visual responses in other classically-defined oculomotor neurons, like brainstem omnipause neurons ^56–62^, would also be expected to exhibit low-pass visual sensory characteristics ^2,63^.

Our results could also help explain how the SC can rapidly detect peripheral object images ^64^ and faces ^65–69^. Indeed, spatial frequency is an important aspect of face and object recognition ^70–74^, but recognition may be thought of as a step following detection. Thus, recognition, which relies more on more band-pass frequencies ^71,72,74^, could benefit from the broader spectrum of V1 preferences that we saw, whereas rapid peripheral object detection could benefit from the SC’s visual properties ^1^. This implies that coarse-to-fine image processing makes most sense in a naturalistic active vision context because it is sufficient to drive rapid foveating eye movements based on coarse peripheral previews.

In this regard, it is worth contemplating our observation of a scarcity, and impoverished representation, of ∼2 cpd preferences in the SC (Fig. 6). The fact that we observed this scarcity and shallow tuning both here and in our earlier study ^15^ makes us believe that it is a genuine property of the SC’s spatial frequency tuning properties. One possibility could be that this scarcity of ∼2 cpd representation is due to the larger SC RF’s than in V1 (Fig. S2B, C). However, this argument is not entirely valid, since it would not explain that we found more SC neurons preferring an even higher spatial frequency (4 cpd) in both studies (and with narrower tuning curves; Fig. 6C). Interestingly, in the SC motor bursts ^38^, 2 cpd spatial frequencies are actually preferred, and we also found here that visual reafferent responses were also elevated for 2 cpd (Fig. S6B). Thus, it would be interesting in the future to find a behavioral correlate of these observations, particularly in the context of active orienting behaviors, and to also map the exact spatial frequency ranges of the underrepresentation seen in Fig. 6.

One should also discuss our V1 observations further. For example, we obtained fundamentally different results from those in V1 of other species ^33,75^. In the more recent study, V1 coarse-to-fine dynamics were clear in both the mouse and marmoset monkey ^33^. However, in that study, the authors analyzed visual reafferent responses after gaze shifts ^33^, whereas we focused on stimulus-evoked responses. In the mouse portion of that study, these authors did relate their active vision observations to stimulus-evoked tuning, finding that the mouse V1 coarse-to-fine dynamics in active vision could be well explained by such tuning during passive viewing. This is very different from both our V1 as well as SC results. In fact, in our SC results, we consistently saw dissociations between visual preference and coarse-to-fine dynamics in terms of visual response latency (e.g. Fig. 1). Thus, there seems to be a fundamental species difference between our observations and those in the mouse and marmoset monkey. This difference, which could stem from the lower importance of V1 in some mammals relative to primates, also persisted in our analyses of visual reafferent responses after microsaccades (Fig. 7).

We also believe that the differences between species extend also to the SC. For example, in a subsequent study ^76^, the same authors also explored coarse-to-fine dynamics in the mouse SC. While they did find a preference for low spatial frequencies, like we did, their observations were again fundamentally different from ours because their mouse V1 and SC findings were more similar to each other than our macaque V1 and SC observations. Also, we saw stronger dissociations between response strength and latency in the macaque SC. It would be interesting in the future to explore species differences in more detail. One possibility could be that we used larger gratings than the V1 RF’s (Fig. S2C; Methods). This was because we sometimes recorded SC and V1 neurons simultaneously, and we matched grating sizes to the larger SC RF’s (Methods and Fig. S2B). Thus, our gratings likely activated the surrounds of V1 RF’s. However, while this is expected to dampen V1 visual responses slightly ^77–80^, it would not entirely eliminate them. Moreover, real-life viewing of natural scenes does not always entail stimuli that are perfectly optimized for V1 neurons. Therefore, in naturalistic settings, our observations on the differences between SC and V1 coarse-to-fine dynamics are likely more realistic than not.

We were also intrigued by the weak V1 coarse-to-fine dynamics that we observed, especially because the substrates for such dynamics are present upstream of the cortex ^26^. Indeed, we decided to look at this issue in our own data, and we did so by classifying neurons according to V1 depth (Methods). We found hints of stronger coarse-to-fine dynamics in a very narrow band of V1 depth (Fig. S7). Besides potentially validating the idea of earlier subcortical substrates for coarse-to-fine processing upstream of V1 ^26^, this suggests that there is a reformatting of inputs within V1 processing itself, just like we argue that there is a reformatting of SC processing from its cortical inputs. Naturally, with even higher spatial frequencies than what we tested, there could be clearer V1 evidence for a delaying of visual responses, like in the SC. However, this would not change the fact that for a broad range of spatial frequencies (like the ones that we considered), such a coarse-to-fine dynamic is much stronger in the SC than in V1.

Finally, the fact that V1 neurons favored slightly more low-pass preferences in visual reafferent responses than in stimulus-evoked activity likely reflects interactions between the orientation of the eye movement vector, its size and speed, and the original grating orientation. This is not too different from what we observed earlier in the SC ^27^.

In all, our results suggest that coarse-to-fine image processing dynamics are much more of a collicular than a cortical visual phenomenon.

## Competing interests statement

The authors declare no competing interests.

## Acknowledgements

We were funded by the Deutsche Forschungsgemeinschaft (DFG; German Research Foundation) through the following projects: SPP 2411 Sensing LOOPS: Cortico-subcortical Interactions for Adaptive Sensing (project numbers: 520617944 and 520283985, HA 6749/11-1); BU4031/1-1; SPP2205 Evolutionary optimization of neuronal processing (project numbers: 430158665 and 430157666, HA6749/3-2); SFB 1233 Robust Vision (project number: 276693517); and BO5681/1-1.

## Methods

### Experimental animals and ethical approvals

Monkeys A and F (male; Macaca mulatta; aged 12 years and 14 years, respectively; weighing 9.5 kg and 13.5 kg, respectively) were used in our experiments. A scleral search coil was implanted into one of the two eyes of each animal, in order to measure eye movements using the magnetic induction technique ^81,82^. The monkeys were also implanted with a head holder, for stabilizing head position, and a recording chamber centered on the midline and angled posterior of vertical by ∼38 deg. This chamber allowed accessing both the SC and V1, as we had done previously ^12,13^.

All experiments were approved by ethical committees at the regional governmental offices of the city of Tübingen (Regierungspräsidium Tübingen).

### Laboratory setup

During the experiments, the monkeys sat in a dim room in front of a ∼30 by ∼23 deg visual angle CRT display. The distance from the animals’ eyes to the display was 72 cm. The display refresh rate was 85 Hz.

The experimental procedure was controlled with a modified version of PLDAPS ^83^, and the stimuli were rendered by the MATLAB Psychophysics Toolbox ^84–86^.

Neurophysiological and behavioral data acquisition was achieved with the OmniPlex system from Plexon (1 kHz sampling rate for the behavioral data and 40 kHz sampling rate for the neurophysiological data). Linear microelectrode arrays (V-Probes) from Plexon were used to record neuronal activity. For monkey A, 16 and 24 channel probes were used in the SC recordings, and 16 channel probes were used in the V1 recordings. For monkey F, 16, 20 and 24 channel probes were used in the SC recordings; 16 and 24 channel probes were used in the V1 recordings.

### Experimental procedures

#### Main visual stimulation paradigm

The monkeys fixated a small white spot (79.9 cd/m^2^, ∼10.8 by ∼10.8 min arc of visual angle) presented over a uniform gray background (26.11 cd/m^2^). After a fixation time of 700 ms to 900 ms, a static Gabor patch with 80% contrast appeared for 300 ms, and then disappeared; the fixation spot remained on the display for another 300 ms, after which the monkeys were rewarded. The location of the stimulus was chosen to cover the receptive fields (RF’s) of both SC and V1 neurons. The size of Gabor patch was 3 deg in radius, and the *σ* parameter of the Gaussian mask was 1.06 deg. The spatial frequency of the Gabor patch varied randomly across trials from among 0.5, 1, 2, 4, and 11 cycles per degree (cpd). Grating phase and orientation were also randomized from trial to trial. Across all experiments, we analyzed 35-333 trials per spatial frequency per session.

The RF’s of the neurons were mapped either with a delayed saccade task or an RF mapping task before running the above paradigm. These tasks are described next.

#### Delayed saccade task

The monkeys maintained fixation on a small white spot (79.9 cd/m^2^, ∼10.8 by ∼10.8 min arc). After a fixation period of 300-700 ms, a target spot spot (79.9 cd/m^2^, ∼10.8 by ∼10.8 min arc) appeared at a random location and stayed on the screen. The monkeys needed to maintain fixation until the fixation spot disappeared, and to then make a saccade to the target location and stay fixated on the target spot for another 500 ms. The time between target spot onset and fixation spot offset varied randomly from 500 ms to 1000 ms. We typically collected 100-200 mapping trials per session.

#### RF mapping task

The monkeys maintained fixation on a small black spot (∼10.8 by ∼10.8 min arc) over the same uniform gray background. After a fixation period of 700-900 ms, a white disk (79.9 cd/cm^2^, 12 min arc radius) appeared for 300 ms and then disappeared for another 300 ms. The monkeys needed to maintain fixation throughout the whole sequence of stimulus onset and offset. We typically collected 100-250 mapping trials per session, sometimes increasing this to 300-400 trials especially when the SC and V1 RF hotspots were not very close to each other.

### Data analysis

We analyzed the activity of 112 SC neurons and 251 V1 neurons, collected across 24 SC sessions and 26 V1 sessions; in 14 of the sessions, both the SC and V1 neurons were recorded simultaneously. The neurons were obtained after offline spike sorting using Kilosort^87^.

We detected saccades and microsaccades using our established methods ^88,89^. We excluded trials with microsaccades occurring within +/-50 ms from stimulus onset, to avoid modulations in visual sensitivity.

To select visually responsive neurons, we performed a left-tailed t-test on the spike counts within the interval 50-150 ms before the stimulus onset and the spike counts within the same interval after the stimulus onset. We only included neurons in our analyses that had significantly higher spike counts after stimulus onset (across all trials regardless of stimulus appearance).

We excluded trials with 11 cpd spatial frequency from further analyses. This was because visual responses were generally weak (and sometimes nonexistent) for this spatial frequency, and the latency of whatever visual response that we got could additionally be influenced by the grating visibility and the contrast sensitivity function (and not just the spatial frequency of the stimulus as per our observations in Results). Therefore, we elected to focus on the spatial frequencies with high visibility. Having said that, inspection of the data did reveal that response latencies for 11 cpd were delayed ^15^ more strongly than in V1, consistent with the rest of the results that we reported here.

#### Visual response latency estimation

We estimated the visual response latency of a given neuron for each spatial frequency separately. We did so by first convolving spike times with an asymmetric firing rate density kernel, as we did recently ^12^, so that we do not blur the visual response latency estimates back in time (had we used a symmetric kernel). Then, we calculated the average firing rate curve across trial repetitions of the same spatial frequency. To estimate the visual response latency from this average firing rate curve, we calculated the maximum of the average curve within the final 150 ms before stimulus onset, and we used this as the threshold. The visual response latency estimate was the first time point after stimulus onset for which the firing rate curve crossed above this baseline threshold.

#### Visual response strength estimation

For all other analyses, we applied a gaussian kernel with standard deviation of 10 ms as our convolution kernel with the spike times. Then, we averaged the firing rate across all trials of a given spatial frequency. The response sensitivity was the peak average firing rate during the interval 0-100 ms after stimulus onset.

#### Additional analyses

To test the effect of brain area on visual response latency, we fitted a linear mixed-effects model with latency as the response variable, brain area and spatial frequency as fixed effects, and neuron identity as random effect. The spatial frequency was modeled as a categorical variable, because we did not assume a linear relationship between response latency and stimulus spatial frequency ^15^.

We also performed pairwise t-tests comparing pairs of visual response strength or latency measurements for pairs of individual spatial frequencies. Specifically, in Fig. 3, we compared 0.5, 2, or 4 cpd responses to responses for 1 cpd. The reported p-values were corrected for multiple comparisons by the Holm–Bonferroni method. In the same figure (Fig. 3), we also measured the difference in responses (between 1 cpd and either 0.5 cpd, 2 cpd, or 4 cpd) as marginal histograms. In each panel, we had two such differences, which we also tested against each other using t-tests.

To summarize our population results, we sometimes calculated an average normalized population firing rate curve. To do this, we first normalized the firing rate of each neuron by the peak firing rate of the neuron’s preferred spatial frequency, and we then averaged the firing rate curves across all neurons with the same spatial frequency preference.

To show the effect of neuron preference on the visual response latency, we fitted linear mixed-effects models with spatial frequency preferences and stimulus spatial frequency as the fixed effects and neuron identity as the random effect. We did this separately for the visual response latencies of the SC and V1.

We also checked visual reafferent responses. We took all microsaccades that happened within 100-250 ms after the stimulus onset. During this period, the stimulus was still on, and the original visual response was already almost fully adapted. Then, we aligned the firing rate data to the microsaccade onset and checked if there was a significant response the eye movements. Specifically, we performed t-tests on the firing rates within 25-75 ms after microsaccades (reafferent responses) to compare them to the firing rates from the trials without microsaccade withing same time window (50 ms) around average microsaccade onset time. We also performed another t-test on the reafferent responses, now comparing them to a baseline -25 ms to 25 ms around the saccade onset. We only took the neurons that passed both tests into the analysis. That is, the activity after microsaccades was considered a genuine reafferent visual response if it was higher than firing rates without microsaccades within the same time period after stimulus onset, and if it was also higher than during the peri-microsaccadic epochs. For population summaries, we normalized according to the peak reafferent response of the preferred spatial frequency.

To summarize spatial frequency tuning curves, we measured visual response as a function of spatial frequency for each neuron. Then, we grouped neurons according to the preferred spatial frequency, and we normalized the firing rate of the population by the condition giving the peak response. We then plotted groups of tuning curves according to preference. For comparing the SC and V1, we aligned the tuning curves to the preferred spatial frequency. In this case, the x-axis became the relative spatial frequency (lower or higher than the preferred spatial frequency).

We also assessed the sizes of the population RF’s across experiments. Since each session had multiple neurons and the same stimulus location, the relative RF positions were jittered in position a bit relative to the stimulus (due to subtle RF position variability from neuron to neuron). To confirm that this jitter was minimal and that our stimuli were relatively centered on the population RF’s, we calculated each neuron’s RF, plotted its surface and normalized it to a peak value (within 3.5 deg from the stimulus location because we anyway targeted the stimulus to be near the RF center positions within each session), then averaged all RF surfaces around the stimulus center. That is, we obtained a population RF relative to the stimulus center. If the stimulus was properly centered across most RF’s, then the population RF should peak near the stimulus center. For estimating the RF surface of each neuron, we measured visual response strength from the RF mapping tasks mentioned above. Then, we interpolated between the measurements to obtain a surface where the x, y values corresponded to stimulus location, and the z value corresponded to neuron response.

Finally, in the SC, we did not separate neurons as being visual or visual-motor, especially because we had already done this before in our earlier study ^15^. That earlier analysis revealed that the same coarse-to-fine dynamics were to be expected from visual and visual-motor neurons (save for a constant difference in visual response latencies between the two neuron types). However, in V1, we did analyze potential influences of cortical depth. Like we did recently ^12^, we used current source density (CSD) analysis ^90^ to align recordings across sessions. To do this, we identified a sustained current sink after stimulus onset ^91^, and we measured the depths of individual neurons from the border between this sustained current sink and a sustained current source right under it. We then repeated our analyses of visual response latencies as a function of spatial frequency, and we plotted these latencies using a moving average of 150 micrometers in steps of 50 micrometers.

## Supplementary information

**Figure S1.**
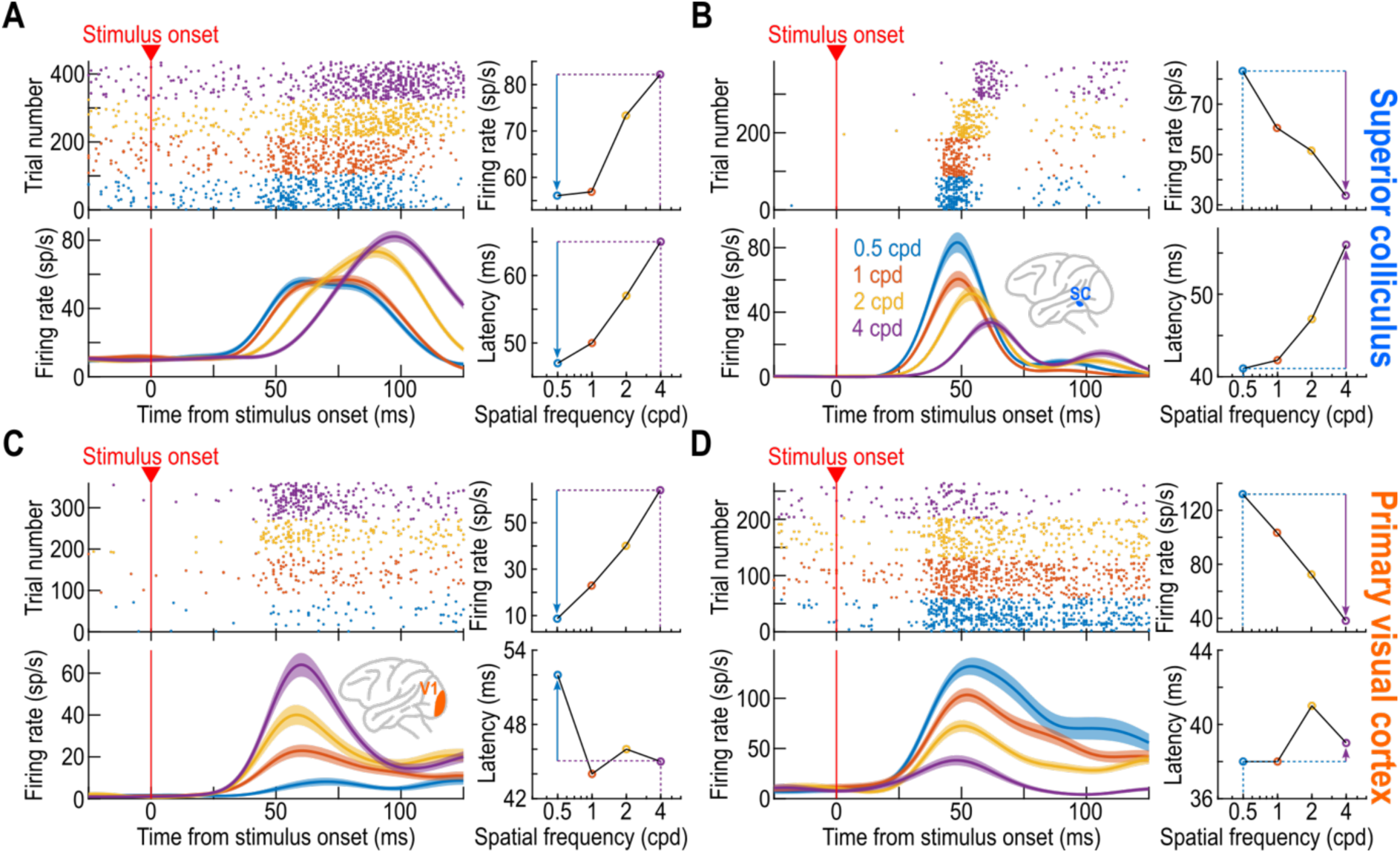
Additional example SC and V1 neurons demonstrating much stronger coarse-to-fine visual processing in the primate SC than in V1. (A) Example SC neuron preferring 4 cpd in terms of visual sensitivity (like in Fig. 1A). The neuron’s visual response latency for lower spatial frequencies was still substantially shorter than its visual response latency for its preferred spatial frequency, and there was a rank ordering of response latencies with increasing frequencies. (B) A second example SC neuron, this time preferring 0.5 cpd. The response latency curve (bottom right) looked similar to that in A, despite the dramatically different sensitivity tuning curves (top right in A, B). (C, D) Example V1 neurons preferring 4 cpd (C) and 0.5 cpd (D) in terms of visual response strength. Here, visual response latency for both neurons was more governed by the sensitivity tuning curve. That is, when the visual response was low (such as at 0.5 cpd for the neuron in C), the visual response latency was longest. The visual response latency for 0.5 cpd was only fastest when this spatial frequency was the preferred spatial frequency of the neuron in terms of its visual response strength (D). All other conventions are like in Fig. 1.

**Figure S2.**
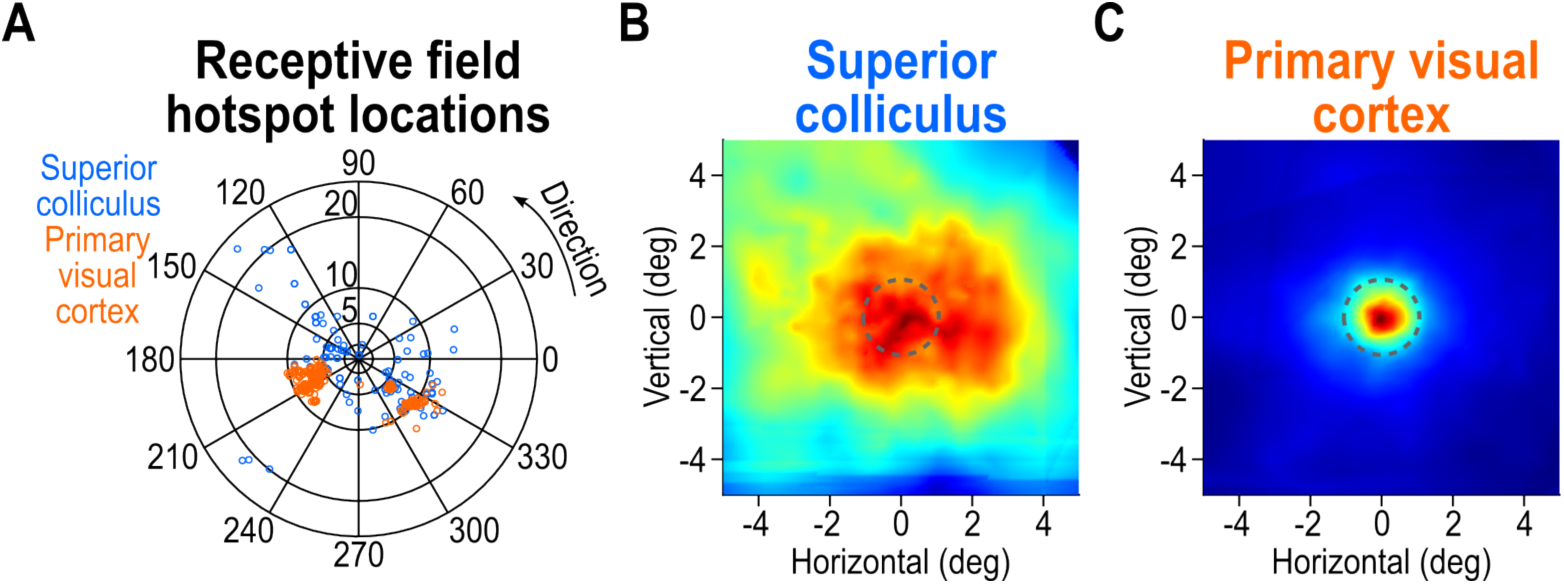
Receptive field (RF) locations and the relationship between RF sizes and our stimuli in the two brain areas. **(A)** Each circle indicates the RF hotspot location of a recorded neuron, from either the SC (blue) or V1 (orange). There was overlap in preferred eccentricities across the collected populations from the two areas, but in the SC, we also sampled the upper visual field (our prior work showed that both the upper and lower visual field representations of the SC, despite their differences ^37,39^, exhibit similar coarse-to-fine dynamics ^15^). **(B)** Average normalized population RF’s of all SC neurons relative to the stimulus location. The origin indicates the center of the stimulus across experiment. The heatmap indicates the average of all normalized SC RF surfaces in our population. Because each electrode track had multiple neurons whose RF centers could be slightly jittered in position relative to each other, the shown population RF slightly overestimates the actual RF sizes of individual neurons. However, the population RF indicates that our stimuli (Methods) were well aligned to RF position, and that they also had sizes within the excitatory portions of the SC RF’s. The dashed circle indicates the *σ* parameter of the Gaussian masks used to generate our Gabor stimuli (overall stimulus radius was 3 deg). **(C)** Average normalized population RF’s of all V1 neurons relative to the stimulus location. Like in **B**, for each neuron, we plotted the normalized RF surface. Then, we averaged all such surfaces around the center of the stimulus across experiments. Our stimuli were well-centered over the V1 RF’s (dashed circle like in **B**). However, they were larger than these RF’s (Methods); nonetheless, we measured robust visual responses (examples are shown in Figs. 1, S1).

**Figure S3.**
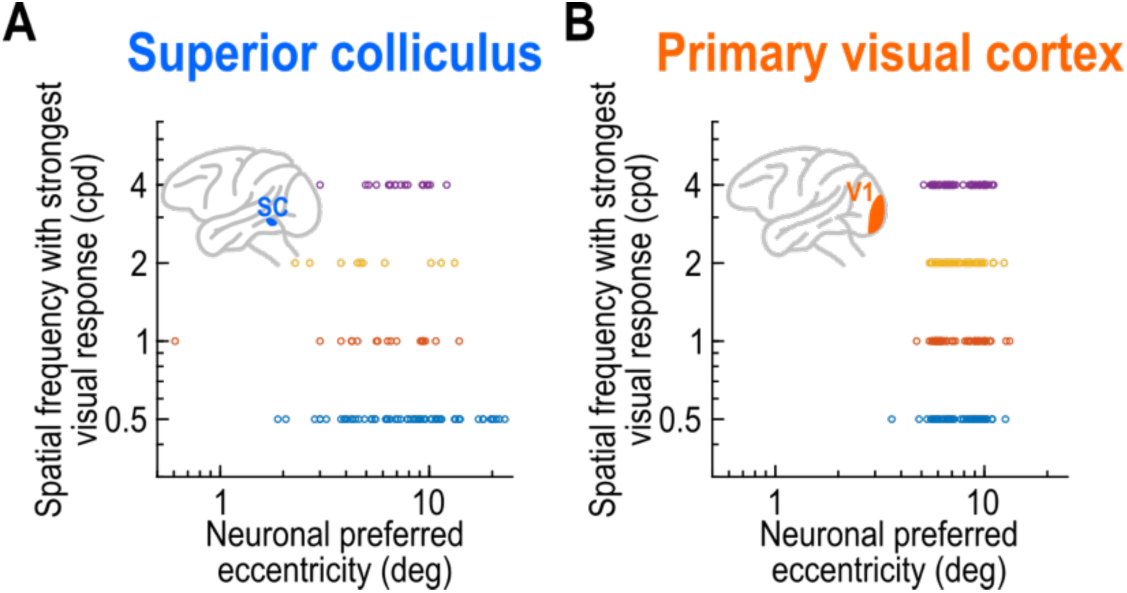
Relationship between preferred RF eccentricity and preferred spatial frequency across our SC and V1 neuronal populations. **(A)** We plotted the distribution in Fig. 6A as a function of the preferred RF eccentricities of the SC neurons (obtained from Fig. S2A). Most neurons preferred the lowest spatial frequency (0.5 cpd), but there were neurons preferring all sampled spatial frequencies. **(B)** Like **A** but for the V1 neurons. All sampled spatial frequencies were preferred equally well across all tested eccentricities.

**Figure S4.**
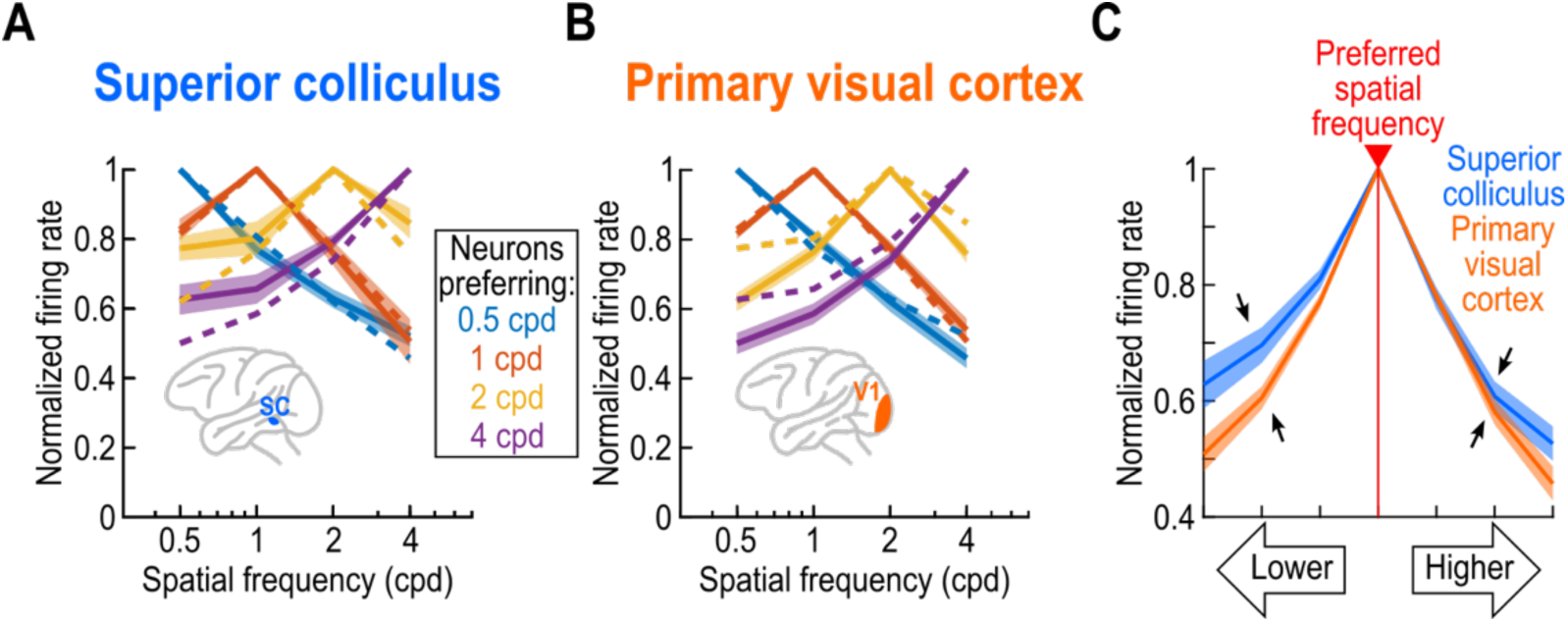
Comparison of the spatial frequency tuning curves in the SC and V1. **(A)** Each curve shows the average spatial frequency curve of all SC neurons preferring a given spatial frequency (like in Fig. 6C), and error bars denote SEM across neurons. The dashed lines show the corresponding V1 tuning curves (from **B**). The SC tuning curves were similar to those in V1, except for two notable differences. For the neurons preferring 2 cpd (yellow), the SC tuning curves were substantially narrower than for the V1 neurons preferring 2 cpd. Second, for the neurons preferring 4 cpd, the lower spatial frequency portion of the neurons’ tuning curves was shallower in the SC than in V1, suggesting that the SC neurons were more sensitive to low spatial frequencies even when they preferred 4 cpd. **(B)** V1 tuning curves (like in Fig. 6D) but overlaid together in the same plot. Now, the dashed curves are the SC tuning curves. **(C)** Within each area, we took each neuron’s preferred spatial frequency and made it the reference spatial frequency (red line). Then, we plotted the neuron’s visual responses for either lower or higher spatial frequencies (ranked in the relative order of our sampled spatial frequencies), and we then averaged across neurons. So, for example, if a neuron preferred 0.5 cpd, then the response at 0.5 cpd was the reference response (red line), and responses to 1, 2, and 4 cpd were plotted at the first, second, and third ordinal positions, respectively, to the right of the red line. Similarly, if a neuron preferred 2 cpd, then the response at 2 cpd was the reference response (red line), and responses to 0.5 and 1 cpd were plotted at the ordinal positions to the left of the red line; and, responses to 4 cpd were plotted at the first ordinal position to the right of the red line. Across neurons, the low spatial frequency portion of the SC tuning curves (blue) was shallower than for the V1 tuning curves, providing further evidence of an emphasis on coarse patterns in the SC. Error bars denote SEM across neurons.

**Figure S5.**
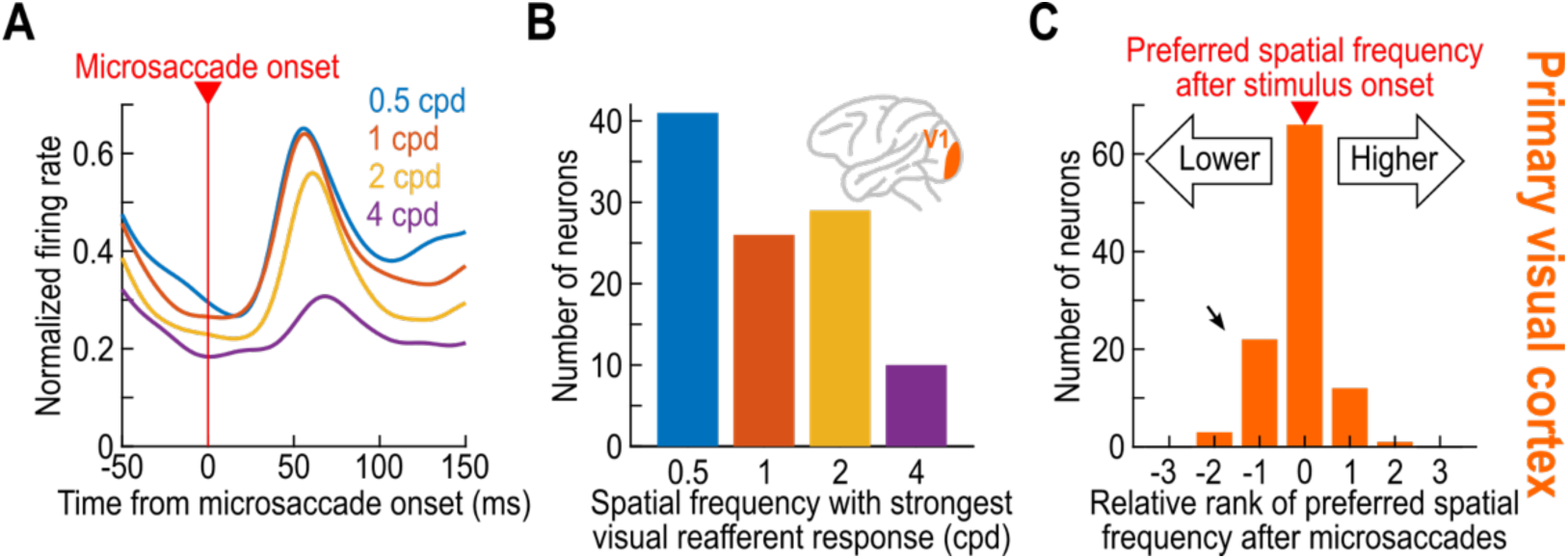
Small shift towards low spatial frequency preference in V1 visual reafferent responses after microsaccades. **(A)** Average normalized population firing rates from all of our V1 neurons when aligned to microsaccade onset. Responses to the lowest spatial frequencies were strongest and fastest when compared to the higher spatial frequencies. **(B)** Distribution of preferred spatial frequencies in our V1 neurons’ visual reafferent responses. There was a clearer dominant preference for low spatial frequencies in the neurons’ postsaccadic visual reafferent responses than in the case of stimulus-evoked responses. **(C)** We compared the preferred spatial frequency in the stimulus-evoked epoch (e.g. Fig. 6B) and the post-saccadic epoch. Zero on the x-axis indicates that a neuron’s preference in the reafferent epoch was the same as in the stimulus-evoked epoch. Neurons at non-zero x-axis values indicate the rank ordering of the shifted preference in the post-saccadic epoch. So, if a neuron preferred 2 cpd in the stimulus-evoked epoch and 1 cpd in the post-saccadic epoch, then it shifted its preference to one lower spatial frequency position in the 4 sampled spatial frequencies. Thus, the neuron was placed in the -1 bin on the x-axis. Across the population, some neurons shifted to preferring lower spatial frequencies in the post-saccadic epoch (oblique arrow), explaining the results of **A**, **B**.

**Figure S6.**
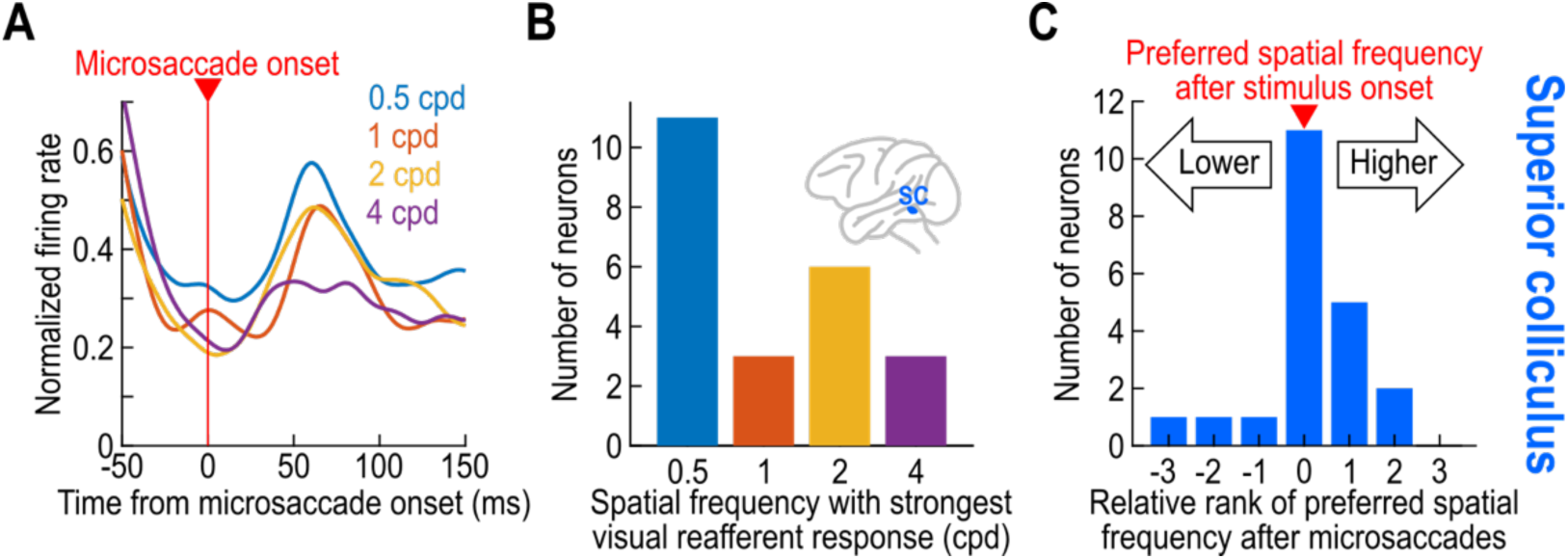
Stronger low spatial frequency preference in SC visual reafferent responses after microsaccades (when compared to V1). **(A)** Similar to Fig. S5A but for the SC neurons. The results replicate our earlier observations from the SC of different monkeys ^27^. **(B)** Similar to Fig. S5B but for our SC neurons. Again, there was a preference for low spatial frequencies. However, note how this effect was stronger than in V1 (see Fig. S5), with most SC neurons preferring 0.5 cpd in their reafferent visual responses. Interestingly, responses for 2 cpd were elevated in this active vision scenario; this is reminiscent of how the SC saccade-related motor bursts also have an increased preference for 2 cpd ^38^. **(C)** Similar to Fig. S5C but for our SC neurons.

**Figure S7.**
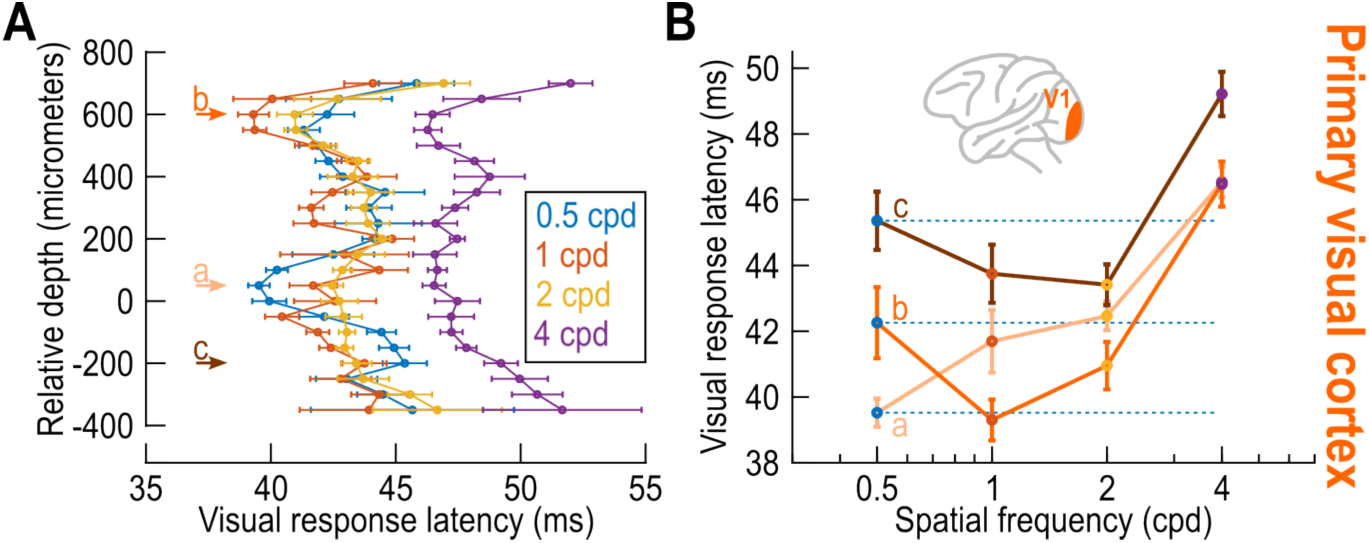
Stronger coarse-to-fine visual processing in a very narrow band of V1 depth. **(A)** We used current source density (CSD) analysis, marking a sustained sink after stimulus onset ^90^ (Methods), to align the relative depths of neurons across all recorded sessions. We then plotted visual response latencies to different spatial frequencies as a function of relative V1 depth. Error bars indicate SEM across neurons within a given bin, and we do not show depth bins for which we had less than 11 neurons in any of the different spatial frequency conditions. Visual response latencies showed two local minima, one at our alignment depth (labeled a) and one above (labeled b). However, only at the alignment depth (labeled a) was there a clear coarse-to-fine temporal dynamic (shorter response latencies for lower spatial frequencies). More superficial (example labeled b) and deeper (example labeled c) V1 neurons did not have the shortest visual response latencies occur for 0.5 cpd. **(B)** Visual response latency as a function of spatial frequency for the V1 neurons at the three example depths shown in **A**. Only within a very narrow band of V1 depths (labeled a) was there a rank ordering of visual response latencies in V1; this could indicate that at this depth, the neurons were more likely to prefer 0.5 cpd spatial frequencies in their visual response strengths.

## Notes

### Competing Interest Statement

The authors have declared no competing interest.

### Summary of Updates

- Added a more detailed V1 depth analysis in Fig. S7 - Added acknowledgements and a competing interests statement - Included minor text corrections

